# Facile real time detection of membrane colocalization of RAS superfamily GTPase proteins in living cells

**DOI:** 10.1101/369587

**Authors:** Yao-Cheng Li, Luke Wang, Tikvah K. Hayes, Margie N. Sutton, Robert C. Bast, Frank McCormick, Channing J. Der, Geoffrey M. Wahl

## Abstract

Members of the RAS family of GTPases (KRAS4A, KRAS4B, HRAS, and NRAS) are the most frequently mutated oncogenes in human cancers. The CAAX motif in the C-terminal hypervariable region (HVR-CAAX domain) contains the cysteine residue that is critical for protein prenylation that enables RAS protein membrane localization, homodimer/oligomer formation, and activation of effector signaling and oncogenic activity. However, it remains unclear if RAS can interact with other prenylated proteins, and if so, how this impacts RAS function. Here we use a newly developed quantifiable Recombinase enhanced Bimolecular Luciferase Complementation strategy (ReBiL2.0) to investigate some of the requirements for RAS superfamily small GTPase protein interactions, and whether this requires cell membrane localization. ReBiL enables such analyses to be done at physiologic expression levels in living cells. Our results confirm that the C-terminal prenylated HVR-CAAX domain is sufficient to direct KRAS and heterologous proteins to colocalize in the cell membrane. We discovered that KRAS also colocalizes with a subset of small GTPase superfamily members including RAC1, RAC2 and DIRAS3 in a prenylation-dependent manner. KRAS colocalization or co-clustering with heterologous proteins can impact KRAS downstream signaling. ReBiL2.0 thus provides a rapid, simple and straightforward method to identify and characterize the colocalization of membrane-associated proteins and to discover agonists and antagonists thereof.

## Results and Discussion

We developed ReBiL as a platform to analyze transient and low-affinity protein-protein interactions (PPIs) in living cells [1]. ReBiL signals depend on both the binding affinity of the interacting proteins and the levels at which they are expressed. Thus, high expression levels of low-affinity PPI’s and low expression levels of high affinity PPI’s could generate similar luminescent signals (Fig. 1a). Although controlled by the same TRE bi-directional promoter, split-luciferase (½luc) fusion-proteins are not always expressed in a 1:1 stoichiometric ratio. These issues can complicate data interpretations. We modified the available ReBiL system (referred to below as ReBiL2.0) to enable precise quantification and comparison of ReBiL luminescent results obtained with different PPI pairs from different ReBiL reporter cells. We normalize luminescent signal with the amount of expressed ½luc fusion protein (explained in detail in Fig. 1 and Methods). We incorporate a single copy of an HA-epitope tag in the linker region of each ½luc fusion protein to enable accurate determination of their expression ratio using a single anti-HA antibody and quantitative western blotting (Fig. 1b). Importantly, as the ReBiL platform is doxycycline inducible, conditions can be established to express ½luc fusion proteins at near-endogenous levels. The ReBiL2.0 system provides a quantitative strategy to significantly enhance the accuracy required to map PPI networks in living cells.

**Figure 1.**
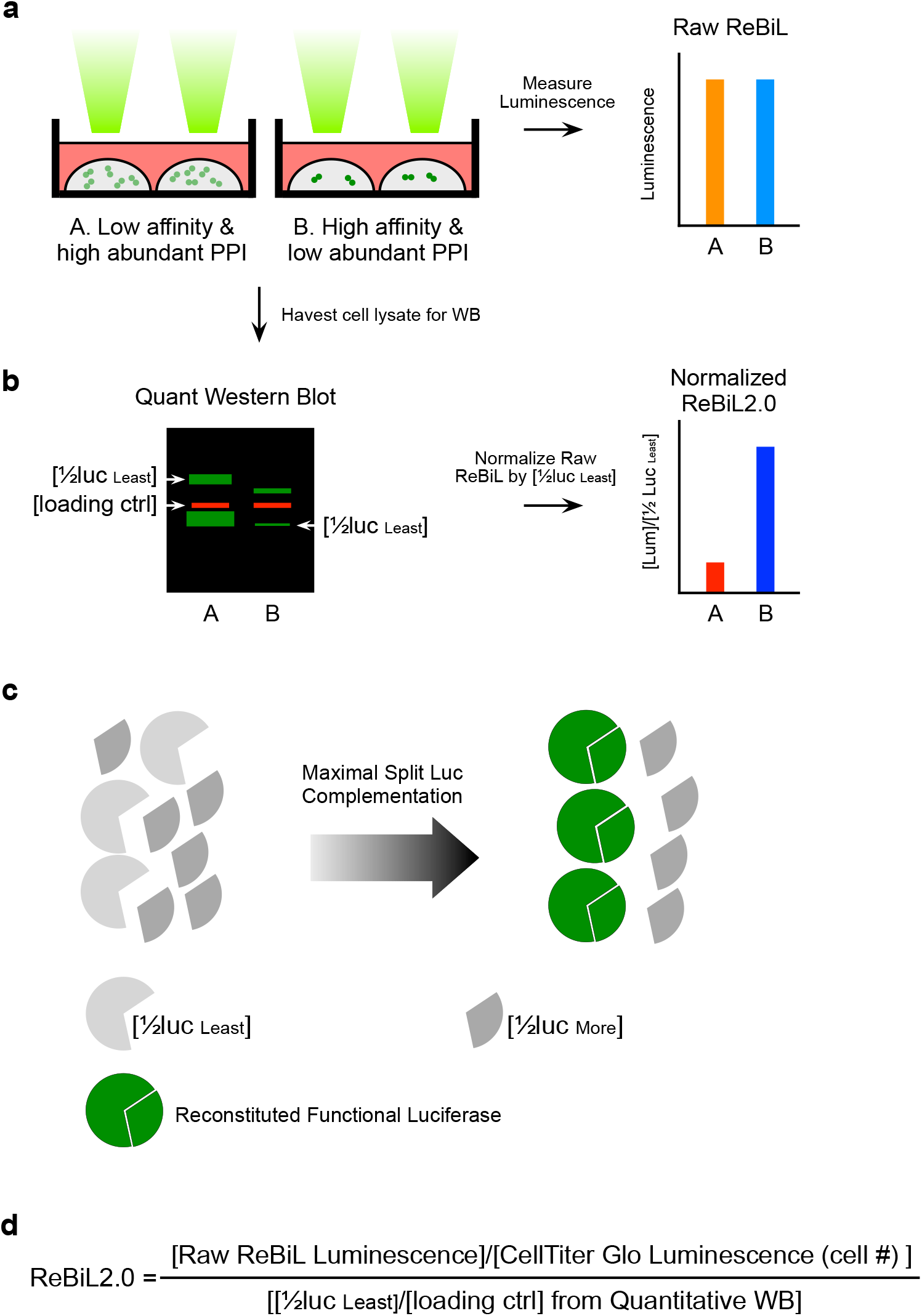
ReBiL 2.0 enables quantification of protein homomer and heteromer formation in living cells. **a,** The raw luminescent signal reported by the original ReBiL assay does not necessarily reflect the affinity of a pair protein-protein interaction since ReBiL signals depend on both the binding affinity of the interacting proteins and the levels at which they are expressed. As low-affinity and high-abundant protein interaction (represented by light green dot pairs) could generate a luminescent signal similar to that resulting from high-affinity and low-abundant protein interaction (represented by dark green dot pairs). **b,** ReBiL2.0 corrects this potential pitfall by normalizing raw luminescent signal to the expressed level of ½luc-fusion. The amount of each ½luc fusion protein and loading control (actin) were quantified by probing western blots with anti-HA and anti-actin antibodies simultaneously using the Licor Odyssey two-channel system (left panel). **c,** Though controlled by the same TRE bi-directional promoter[1], ½luc fusion-proteins are not always expressed in a 1:1 stoichiometric ratio. Thus, the maximal luminescent output generated by reconstituted split-luciferase is determined by the least abundant ½luc fusion [½luc least] when the ½luc fusions are not expressed in 1:1 stoichiometry. Since each ½luc fusion protein contains a single copy of an HA-epitope tag in the linker region (see Fig. 1a), quantitative western blotting with anti-HA antibody enables accurate determination of their relative expression levels as shown in **b**. **d,** ReBiL 2.0 formula. The raw ReBiL signal **[Raw ReBiL Luminescence]** is measured by a luminometer. Then, the relative viable cell number in each sample is measured using CellTiter Glo **[CellTiter Glo Luminescence (Cell #)]** at the end of a kinetic ReBiL2.0 assay. The **[½lucleast]** and **[loading ctrl]** were determined by the corresponding band intensities in western blots using the LI-COR Odyssey system as shown in **b**.

C-terminal HVR-CAAX domain prenylation is required for RAS membrane localization and homodimer or nanocluster formation [2]. Importantly, the prenylated protein database predicts more than 200 prenylated human proteins [3], raising the possibility that RAS proteins might interact with some of them. We used ReBiL2.0 to investigate if KRAS4B (referred to as KRAS unless stated otherwise) can colocalize or co-cluster with other prenylated proteins in living cells, and we then investigated the impact of such colocalization on downstream signaling in the MAPK pathway. To validate this assay, we first asked if KRAS homo-oligomers or nanoclusters are generated in engineered KRAS ReBiL cells (Fig 2a). We appended N-terminal split luciferase (**nl**) and C-terminal split luciferase (**cl**) [1] to the N-terminus of KRAS (designated nl-KRAS and cl-KRAS respectively, Fig 2a). This fusion configuration maintains the intact C-terminal HVR-CAAX domain required for KRAS prenylation. These fusions did not affect KRAS activity as determined by NIH3T3 growth assays (Fig. 2b). When expressed at physiologically relevant levels (Fig. 2c), nl-KRAS and cl-KRAS generated robust luminescent signals (Fig. 2d). Since reconstitution of a functional split luciferase requires a minimum 1:1 ratio of nl-KRAS and cl-KRAS, we interpret these results to indicate that KRAS can form at least homodimers, which is consistent with prior observations [4]. However, formation of KRAS oligomers or nanoclusters [5] cannot be excluded.

**Figure 2.**
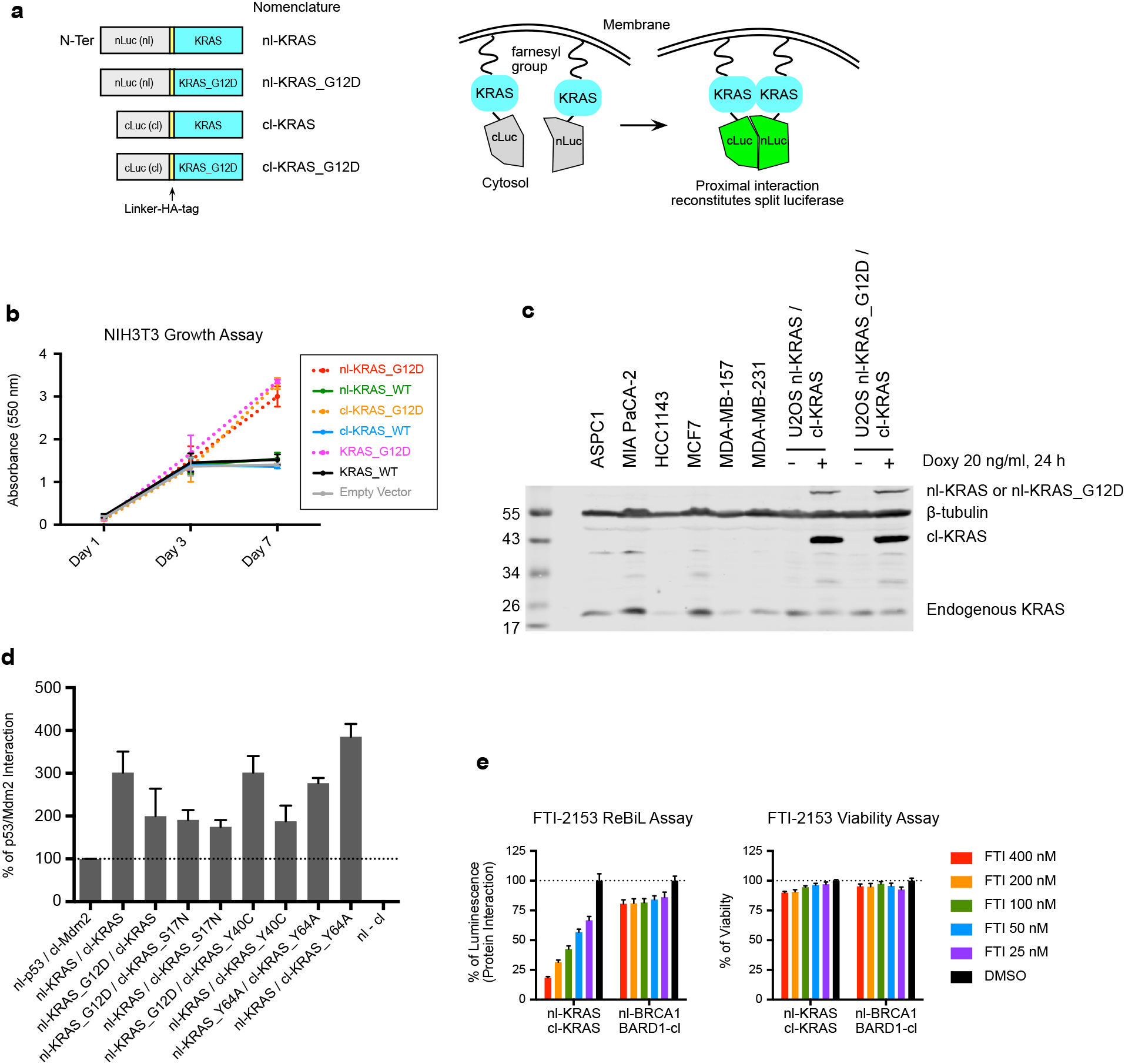
ReBiL 2.0 shows KRAS dimer/oligomer formation unaffected by G-domain mutations. **a,** Diagram depicts the configuration and nomenclature of split-luciferase (½luc) KRAS fusion proteins. The N-terminal ½luc (nl) or C-terminal ½luc (cl) followed by a linker and HA-epitope tag were fused to the N-terminus of KRAS to preserve the integrity and function of the C-terminal HVR-CAAX domain. Right panel illustrates that membrane-associated proximal interaction of the ½luc KRAS fusion proteins (nl-KRAS/cl-KRAS) reconstitutes functional luciferase to generate luminescent signals. **b,** The growth of NIH3T3 cells infected by retroviral vectors encoding ½luc-KRAS fusions or unmodified KRAS was measured by MTT assays at day1, day3, and day7. **c,** Western blot shows that the endogenous KRAS levels in a panel of human cancer cells relative to that achieved by doxycycline (doxy, 20 ng/ml) induction of nl-KRAS/cl-KRAS, nl-KRAS_G12D/cl-KRAS in ReBiL cells. The ½luc-KRAS fusion proteins and the endogenous KRAS were probed by anti-KRAS (mouse monoclonal, Sigma, WH0003845M1) antibody and β-tubulin serves as a loading control. **d**, ReBiL 2.0 shows the quantitative homo/heteromer formations between ½luc KRAS fusion proteins carrying different mutations in their G-domains. The bar chart reports quantitative protein interactions normalized to the well-established nl-p53/cl-Mdm2 (positive control, set to 100%) and the background luminescence generated by ½luc alone lacking fusion partners (nl/cl, negative control, set to 0%) (see Methods). Data show mean ± SEM (n ≥ 3). **e,** Farnesyltransferase inhibitor (FTI-2153) dose-dependently decreased luminescent signals in cells expressing nl-KRAS/cl-KRAS. ReBiL cells carrying nl-KRAS/cl-KRAS were treated with FTI-2153 and Doxycycline. The nl-BRCA1 and BARD1-cl ReBiL cells[1] serve as a control for non-specific effects of the FTI-2153. End-point luciferase and cell viability assays using One-Glo and CellTiter-Glo respectively were performed 24 hours after ReBiL cells were treated with inhibitor (see Methods). Representative data from several biological repeats. Data shown here represent the mean ± SD from eight technical repeats.

We next determined if KRAS activation or effector interaction are required for dimer/oligomer (hereafter referred to as multimer) formation. KRAS_G12D and KRAS_S17N mutants interact with wild type KRAS_WT (Fig. 2d), indicating that the GTP-bound active form (represented by the KRAS_G12D mutant) and GDP-bound inactive form (represented by the KRAS_S17N mutant) have no impact on multimer formation. In addition, the effector binding mutant KRAS_Y40C [6] interacts well with KRAS_WT, indicating that Raf effector binding is dispensable for KRAS multimer formation [4]. Although Tyr64 has been reported to be critical for HRAS dimerization in artificial membranes [7], we observed no measurable effect of a Y64A mutation on KRAS dimer formation in living cells (Fig 2d) [4] even though HRAS and KRAS share over 90% sequence identity in the G-domain [2] including Tyr64. Taken together, these data are consistent with previous results showing that the G-domain plays a minimal role in KRAS dimer/oligomer formation, and they validate the use of ReBiL to investigate Ras homo- and heteromeric interactions.

Previous studies showed that a single Cys to Ser mutation (KRAS_C185S) in the CAAX motif farnesylation site both abolishes RAS protein membrane association [8] and inactivates downstream signaling [4]. Indeed, the farnesyltransferase (FTase) inhibitor FTI-2153 [9] dose-dependently reduced the luminescent signal generated by nl-KRAS/cl-KRAS complementation (Fig. 2e). As FTI treatment affected neither the luminescence generated by BRCA1-BARD1 RING-domain [1], nor viability (Fig. 2e), we conclude that the ability of the inhibitor to reduce the luminescent signal from nl-KRAS/cl-KRAS complementation is not due to an indirect effect of FTI-2153, but rather to its ability to inhibit farnesyltransferase and to subsequently block KRAS farnesylation and membrane localization. This is a surprising result because it has been shown that KRAS can be prenylated by geranylgeranyltransferase I (GGTase I) when FTase is inhibited by FTI in cells that overexpress pMEV [10,11], a mutant transporter protein that facilitates mevalonate uptake [12]. This alternative prenylation restores KRAS membrane localization and function that results in resistance of FTI inhibition [2]. However, the alternative prenylation process might not be efficient in U2OS cells as treatment of U2OS with an FTI has previously been reported to prevent KRAS prenylation [13]. Thus, we interpret our results to indicate that the FTI mediated reduction in nl-KRAS/cl-KRAS association derives from its ability to prevent KRAS membrane association. These results suggest that in addition to HRAS mutant cancers, FTI could have beneficial effects in those KRAS mutant cancers where GGTase I does not contribute significantly to KRAS modification in the context of FTI treatment. While drugging of mutant RAS in cancers has been largely unsuccessful [14], we suggest that the KRAS ReBiL strategy could provide a rapid, high throughput platform for identifying therapeutics that inhibit RAS membrane localization, multimerization, and activity in specific cancers.

As expected, a single cysteine mutation in nl-KRAS_C185S abolishes luciferase complementation (Fig. 3b), consistent with the notion that HVR-CAAX farnesylation is critical for KRAS membrane localization and multimer formation [4]. We next determined if CAAX domains from HRAS and NRAS enable heteromer formation with KRAS. To focus on the HVR-CAAX domain, we replaced the G-domains of each RAS isoform with the monomeric blue fluorescent protein mTagBFP (tBFP) [15]. The ½luc and tBFP fusions are designated nl-tBFP-HVR-CAAX_H_, nl-tBFP-HVR-CAAX_N_, and nl-tBFP-HVR-CAAX_K_ to indicate the origins of HVR-CAAX domains from H, N and KRAS, respectively (Fig. 3a). The nl-tBFP-HVR-CAAX_N_, and nl-tBFP-HVR-CAAX_K_ efficiently reconstituted luciferase activity efficiently when paired with cl-KRAS (Fig. 3b), indicating that KRAS forms heteromers with NRAS in a G-domain independent fashion. By contrast, we consistently observed that nl-tBFP-HVR-CAAX_H_ and cl-KRAS interact less efficiently than nl-KRAS/cl-KRAS and nl-tBFP-HVR-CAAX_K_/cl-KRAS pairs (Fig. 3b). This indicates that HRAS has a lower probability of interacting with KRAS, which is consistent with a previous study showing that KRAS and HRAS can be co-clustered through lipid-mediated spatial cross talk [5]. Consistent with a previous study [4], the CAAX domain itself appears to be sufficient for driving proximal membrane localization since appending only nl and cl to the HVR-CAAXK (nl-HVR-CAAX_K_ and cl-HVR-CAAX_K_) enabled efficient complementation, while nl/cl alone and cl-KRAS/nl-tBFP (without HVR-CAAX) exhibited negligible luminescent signals (Fig. 3b). Thus, we refer to these G-domain independent, but HVR-CAAX dependent, interactions as "proximity enabled" or “proximal” interactions or protein “co-clusters”. We use the latter term as physical measurements suggest that RAS can localize in the membrane as multimeric structures [5]. Importantly, the KRAS proximal interactions are not significantly influenced by complementation of the two ½luc moieties alone, as the C185S mutant did not generate an appreciable luminescent signal (Fig 3b).

**Figure 3.**
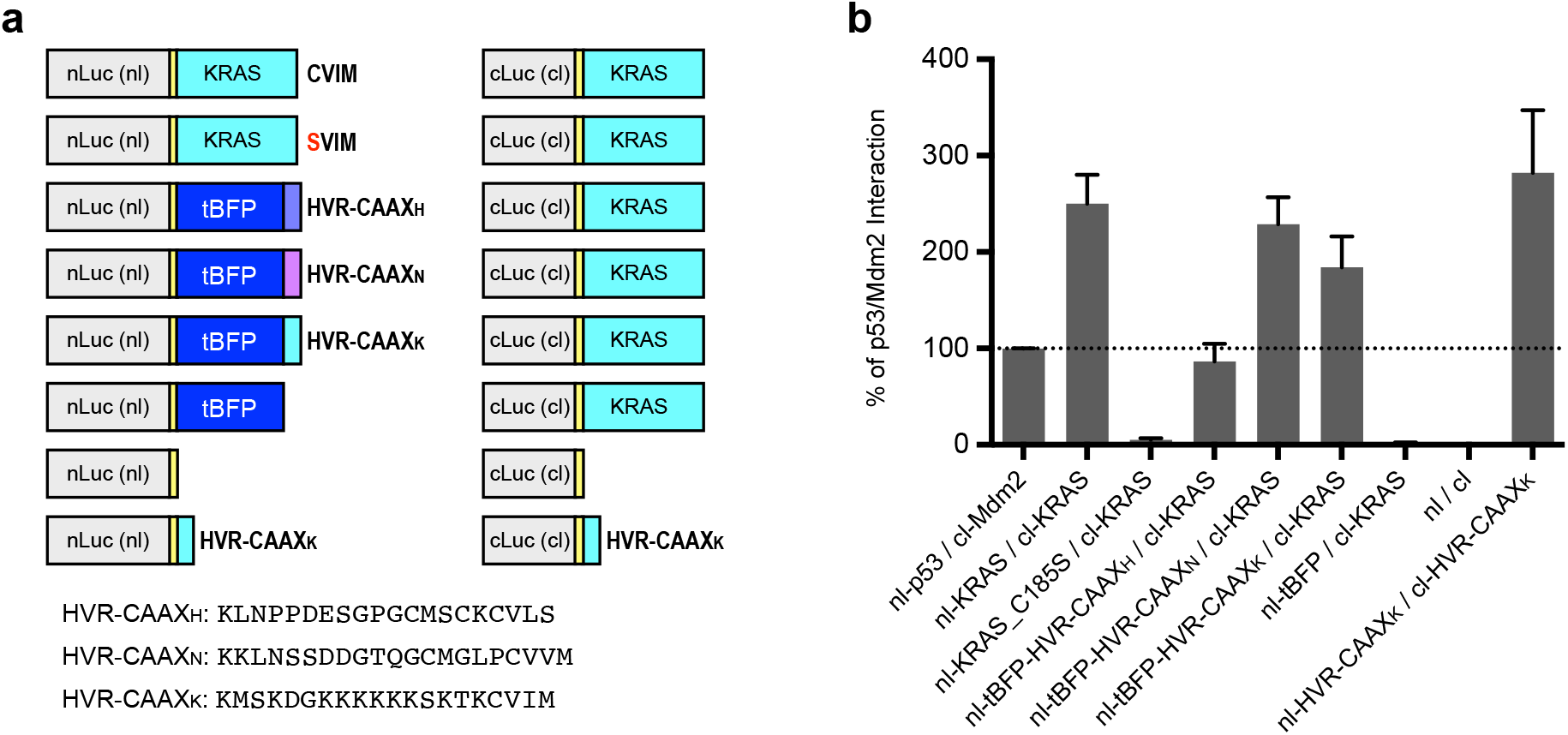
Prenylation on the HVR-CAAX domain is sufficient to generate an area code targeting RAS to specific membrane domains enabling proximal interactions. **a,** Diagrams depict the ReBiL pairs used here. The nl-TagBFP (nl-tBFP) was fused with C-terminal from HRAS, NRAS, and KRAS4B (nl-tBFP-HVR-CAAXH, nl-tBFP-HVR-CAAXN, and nl-tBFP-HVR-CAAXK respectively). The amino acid sequences of three HVR-CAAX isoforms were also shown. **b,** The quantitative homo/heteromer formation between cl-KRAS and nl-tBFP appended to different HVR-CAAX domains are reported after normalization to nl-p53/cl-Mdm2 (100%) and nl/cl (0%). Data represent the mean ± SEM (n ≥ 3).

The above results establish the ability of the ReBiL2.0 assay to capture interactions between prenylated proteins located in membrane domains that coincide with or overlap those occupied by KRAS. Interestingly, KRAS4A, which has a different HVR-CAAX domain [16] than the KRAS4B splice variant, produced a significant ReBiL signal with KRAS4B (Fig 4a), implying that KRAS4A and KRAS4B reside in overlapping membrane domains. These results suggest that KRAS4A and 4B might exhibit functional similarities that could impact each other’s downstream effects [16].

**Figure 4.**
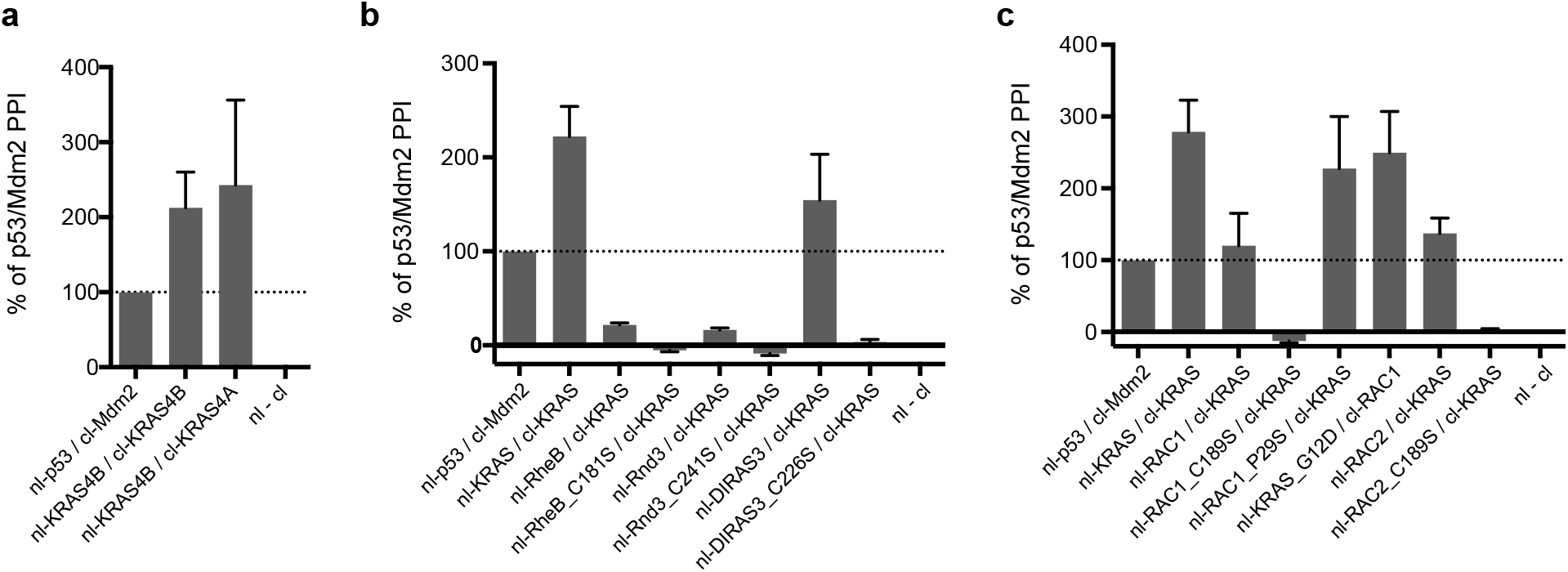
KRAS forms heteromers with other prenylated proteins. **a,** ReBiL 2.0 reveals interaction between KRAS4A and KRAS4B. Data are mean ± SEM (n=2). **b,** ReBiL 2.0 shows KRAS interacts with DIRAS3 in a CAAX-dependent manner, but does not interact with RHEB and RND3. Data show the mean ± SEM (n ≥ 3). **c**, Proximal interaction between KRAS/RAC1 and KRAS/RAC2 depend on an intact CAAX motif. G-domain mutations such as KRAS_G12D and RAC1_P29S do not affect their interaction. Data show mean ± SEM (n ≥ 3).

These observations led us to wonder whether other prenylated proteins are sufficiently near the membrane domains in which KRAS resides to enable heteromeric interactions. As a specificity control, Fig. 4b shows that KRAS does not interact with RheB, a prenylated protein localized to endomembranes [17], not the plasma membrane in which RAS proteins reside. KRAS also does not interact with Rnd3 (Fig. 4b), which also localized to the plasma membrane [17]. This suggests that they reside in sufficiently different plasma membrane domains as to preclude interaction. In contrast, we discovered that KRAS does interact with the genetically defined KRAS suppressor DIRAS3 (Fig. 4b, and Sutton et al, in revision), RAC1 and RAC2 (Fig. 4c) in a prenylation-dependent fashion since a Cys to Ser mutation in the CAAX motif of each protein abolishes luciferase complementation (Fig. 4b, c). The functional importance of such interactions is indicated by DIRAS3, which is a tumor suppressor that downregulates MAPK activity [18] by a mechanism facilitated by its membrane colocalization with RAS proteins (Sutton et al, in revision).

Chemically-induced KRAS_G12D homodimerization can activate the MAPK pathway [4]. Consistent with this, we observed that doxycycline-induced expression of nl-KRAS_G12D and cl-KRAS_G12D also activates the MAPK pathway as indicated by increased phosphorylation of AKT and ERK1/2 (Fig. 5a, b) in ReBiL cells. However, we wondered about the extent to which association between the ½luc moieties could contribute to protein interactions in general, and to MAPK signaling via the RAS pathway in specific. First, we did not detect ½luc (nl/cl) complementation in the absence of the RAS HVR-CAAX domain (Fig. 3b). Second, we previously showed that preformed p53-Mdm2 complexes (p53-Mdm2 Kd = 600 ~ 420 nM [19]) appended with split luciferase complementing pairs can be readily and dose dependently disrupted by the small molecule Nutlin-3a [1]. This indicates that 1/2luc association using the constructs employed in the ReBiL system must contribute little to protein-protein interactions in this system. By contrast, Nutlin-3a did not disrupt p53-Mdm2 complexes when each protein was fused to split NanoLuc [20]. These data reveal that complexes formed with split NanoLuc molecules are very stable, as has been observed with split green fluorescent protein (GFP) complementation (dissociation rate estimated as ~ 9.8 years [21]). Third, we previously showed that the ReBiL assay successfully detects the weak and transient interactions between the E3 ubiquitin ligase FANCL and Ube2t, its cognate E2 ubiquitin conjugating enzyme. By contrast, only background luminescent signal was detected when assaying interaction between a FANCL RING domain mutation (FANCL_C307A) and Ube2t [1]. These results indicate that the affinity between the two ½luc moieties must be smaller than that of FANCL-Ube2t. Together, these data are consistent with the two firefly ½luc moieties having very low affinity for each other.

**Figure 5.**
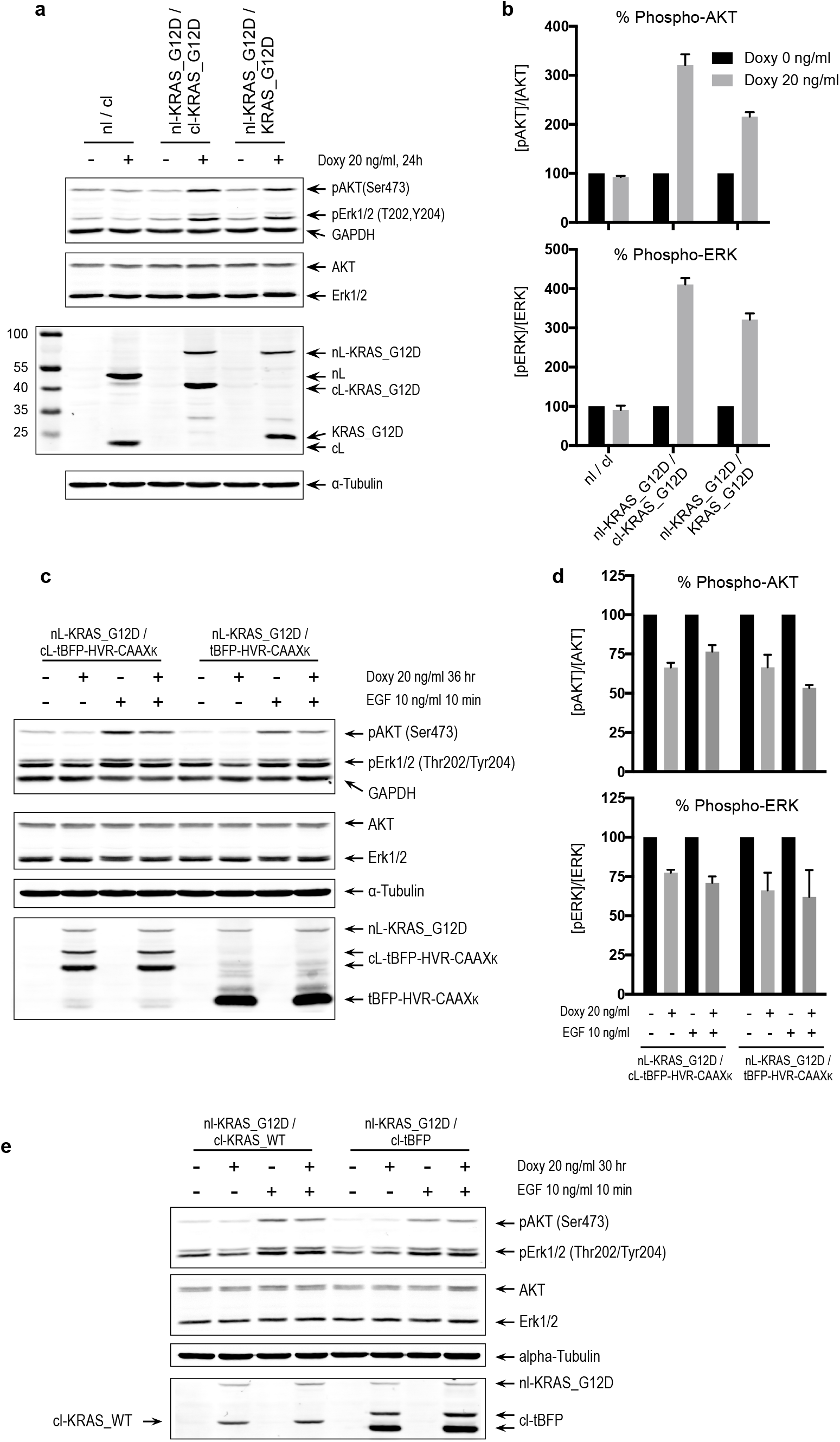
Co-clustering of prenylated proteins can modulate KRAS downstream signaling. **a,** A representative western blot shows that the nl-KRAS_G12D/cl-KRAS_G12D and nl-KRAS_G12D/KRAS_G12D (without fusion to cl) ReBiL pairs activate AKT and ERK1/2 phosphorylation in response to doxycycline induction. The nl/cl pair serves as the negative control. **b,** Quantitation of phospho-ATK (top panel) and phospho-ERK1/2 signals (bottom panel) from (**a**). The amount of phospho-protein from cells without doxycycline (doxy 0 ng/ml) was set to 100% (see Methods). Data represent the mean ± SEM (n=4). **c,** A representative western blot shows that the cl-tBFP-HVR-CAAX_K_ and tBFP-HVR-CAAX_K_ (without fusion to cl) attenuate phosphorylation of ATK and ERK1/2 activated by doxycycline-induced nl-KRAS_G12D in serum-free medium followed with or without EGF stimulation. **d,** Quantitation of phospho-ATK (top panel) and phospho-ERK1/2 (bottom panel) data from (**c**). The amount of phospho-protein from cells without doxycycline (doxy 0 ng/ml) was set to 100%. (see Methods). Data represent the mean ± SEM (n=3). **e,** A western blot shows that the cl-KRAS_WT can weakly attenuate phosphorylation of ERK1/2 in the presence of doxycycline-induced nl-KRAS_G12D in serum-free medium without EGF stimulation. The cl-tBFP (without HVR-CAAX domain) serves as the control.

While the contribution to protein association is likely negligible, it must be non-zero as the ½luc moieties must associate to some degree to reconstitute, if transiently, a functional protein. These results indicate that interactions enabled by proximity of proteins within membrane domains play a dominant role in regulating KRAS signal output. The data in Fig. 5 a and b indicate that less than 20-30% of the effect on KRAS signaling is attributable to split luciferase association. Computational and biochemical studies suggested that the affinities of KRAS dimers are in the order of millimolar (5 mM - 107 mM) [22]. Our results thus suggest that the affinity of split luciferase moieties should be weaker than that. Such weak binding affinity likely could not stably support enzymatic reactions unless they were locally concentrated by, for example, the membrane anchoring enabled by the CAAX domains in nl-CAAXK/cl-CAAXK fusions (Fig 3b).

We next determined whether proximal interactions between KRAS and a nonfunctional partner could attenuate its activity. To this end, we used the cl-tBFP-HVR-CAAXK because it colocalizes with KRAS but lacks a functional G-domain. The cl-tBFP-HVR-CAAX_K_ reduces both phospho-AKT and phospho-ERK1/2 signals in the presence of serum starvation and EGF stimulation (Fig. 5c, d). These results indicate that KRAS activity can be impacted by proximal interactions with heterologous proteins even if they lack appreciable interaction affinity. Similar results were also observed when the cLuc (cl) was removed from cl-tBFP-HVR-CAAXK (Fig. 5c, d), indicating that the observed effect did not derive from low affinity ½luc interactions. These data are consistent with prior reports that a wild-type RAS allele acts as a suppressor of mutant RAS in murine model studies [23–25]. We confirmed this finding in the ReBiL system (Fig. 5e), although the degree of inhibition was greater using tBFP than WT RAS. We reason that the cl-tBFP-HVR-CAAX_K_ inhibits KRAS_G12D better than WT RAS (Fig. 5c, d) because it lacks a functional G-domain, enabling more significant interference with the function of the KRAS mutant and an inability to be cross-activated through KRAS_G12D-mediated recruitment of a guanine nucleotide exchange factor (e.g. SOS1) [26]. Taken together, our results suggest that KRAS signaling output can be fine-tuned by proximity mediated interaction of homologous or heterologous proteins.

RAS activity can be inhibited by the Ras binding domain (RBD; amino acid 51-220) of cRaf (or Raf1) [27]. Since the results presented above emphasize the importance of membrane mediated proximal interactions or co-clustering to regulate RAS activity, we hypothesize that targeting the RBD to the membrane should inhibit RAS more potently through a combination of proximity facilitated interaction or generation of RBD-RAS co-clusters added to the affinity of the RBD for RAS. To this end, we fused the HVR-CAAXK to the C-terminus of RBD. We added to the RBD a destabilizing domain (DD, i.e., FKBP12_F36V_L106P) to enable regulation of protein levels [28] (designated TRE-DD-RBD-CAAXK). As a negative control, we generated TRE-DD-RBD, which should not co-cluster with RAS (Fig. 6a). Both constructs were integrated as a single copy into the U2OS parental ReBiL cell line. While the small molecule ligand Shield1 was not effective in these DD-fused constructs (data not shown), the results clearly show that DD-RBD-CAAXK inhibited both Erk1/2 phosphorylation (Fig. 6b) and cell proliferation (Fig. 6c) far better than similar levels of DD-RBD. These results are reminiscent of an early study showing that despite high binding affinity (6.2 nM to RAS(GTPγS)), only a membrane associated intrabody (iDab#6-memb) exhibits significant inhibitory effects in vivo [29]. However, the importance of membrane co-clustering of Ras with the intrabody was not discussed in that study [29].

**Figure 6.**
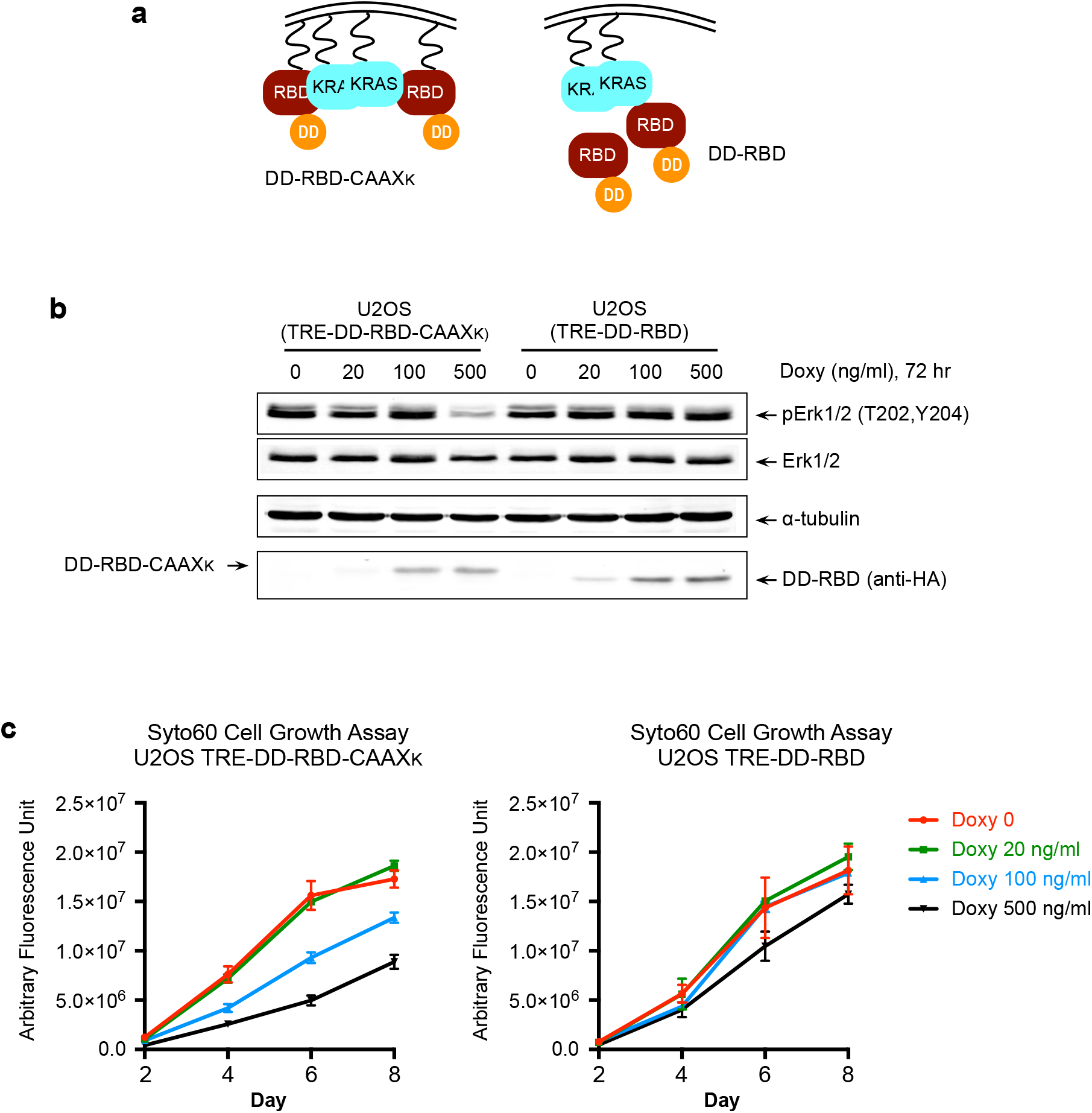
HVR-CAAXK fused RBD shows greater RAS inhibition. **a,** Diagram illustrates the proposed model of action of DD-RBD-CAAXK and DD-RBD. **b,** A representative western blot shows that the DD-RBD-CAAXK inhibits ERK1/2 phosphorylation in response to doxycycline induction whereas DD-RBD lacks inhibitory effect. **c,** Syto60 cell growth assay shows that DD-RBD-CAAXK shows inhibits cell growth in a doxycycline concentration dependent manner. The control DD-RBD shows a very minor inhibitory effect.

In summary, our results reveal that the ReBiL2.0 assay accurately captures KRAS membrane spatial organizations studied by diverse methods including photoactivated localization microscopy (PALM) [4], electron microscopy (EM) [30] and other imaging methods (reviewed in [5]). Our results validate the ReBiL2.0 assay as a robust tool to study membrane-anchored proteins and their interactions, and to identify new interaction partners in living cells. Indeed, we discovered that KRAS4B can colocalize with KRAS4A as well as with a subset of small GTPase family proteins such as RAC1, RAC2, and DIRAS3. We demonstrate its use for studying structure-function requirements for such interactions, and for documenting the mechanisms by which DIRAS3 interacts with and suppresses KRAS function (Sutton et al, in revision). Our results emphasize the importance of proximity enabled co-clustering of KRAS_G12D with wild type RAS and other proteins as homo and heteromers to modulate downstream activity [23–25]. Our results raise possibility that colocalization or co-clustering of membrane proteins may enable orchestration of diverse environmental cues to optimize cell fitness. The existence of hundreds of lipidated membrane proteins and their importance in cellular signaling and tumorigenesis, exemplified by KRAS, emphasizes the need to map and characterize these previously unexpected co-clustered proteins. The ReBiL2.0 platform is an economical, portable and straightforward method which provides a robust and unbiased strategy to identify networks of co-clustered proteins, and to screen for drugs that modulate their activities.

## Acknowledgments

We thank M. Philips, N. Rosen, Z. Yao, M. McMahon, and E. Stites for constructive comments. Work in the laboratory of G.M.W. was supported, in part, by the Cancer Center Core Grant (CA014195), the Susan G. Komen Foundation (SAC110036), the Leona M. and Harry B. Helmsley Charitable Trust (2012-PG-MED002), the Freeberg Foundation, the Greenfields, Sorrento Biosciences and Genentech.

## Author Contributions

Conceptualization, Y.-C.L. and G.M.W.; Methodology, Y.-C.L. and G.M.W., Investigation, Y.-C.L., L.W., and T.K.H.; Writing-Original Draft, Y.-C.L. and G.M.W.; Writing-Review & Editing, Y.-C.L., G.M.W., L.W., M.N.S., R.C.B. C.J.D., and F.M.; Funding Acquisition, G.M.W.; Resources, M.N.S, R.C.B., C.J.D., and F.M.

## Author Information

Correspondence and requests for materials should be addressed to G.M.W.

The authors declare no competing financial interests.

## METHODS

### Construction of ReBiL targeting plasmids and cell lines

The methods for constructing ReBiL plasmids and cell lines have been previously described [1]. ReBiL cell lines were maintained in Dulbecco’s modified Eagle’s medium (DMEM, Corning 10-013-CV) with 10% (v/v) fetal bovine serum (FBS), 10 μg/ml ciprofloxacin (Corning 61-277-RG), 200~400 μg/ml G418 (Corning 61-234-RG), 1 ng/ml doxycycline (Sigma), and 4 μg/ml blasticidin (Thermo Fisher). All ReBiL targeting plasmids and cell lines used in this report are listed in Supplemental Table.

### Antibodies

Antibodies used here are anti-KRAS (mouse monoclonal, Sigma, WH0003845M1), anti-β-tubulin (rabbit polyclonal, LI-COR, P/N 926-42211), anti-HA-tag (rabbit monoclonal, Cell Signaling, #3724), anti-β-actin (mouse monoclonal, LI-COR, P/N 926-42212), anti-phospho-AKT (rabbit monoclonal, Cell Signaling, #4060), anti-AKT (mouse monoclonal, Cell Signaling, #2920), anti-phospho-ERK1/2 (rabbit monoclonal, Cell Signaling, #4370), anti-ERK1/2 (mouse monoclonal, Cell Signaling, #4696), anti-GAPDH (rabbit polyclonal, Santa Cruz, sc-25778), anti-α-tubulin (mouse monoclonal, Sigma), goat anti-rabbit secondary antibody (Alexa Fluor 680, Thermo Fisher, A-21109), and goat anti-mouse secondary antibody (IRDye 800CW, LI-COR, P/N 926-32210).

### ReBiL2.0 assay

This assay consists of **(a)** real-time BiLC assay that has been described previously[1] and **(b)** quantitative western blotting to determine the amount of each ½luc fusion protein. The assay medium (I) consists of DMEM/F12 (Thermo Fisher, phenol-red free), 10% (v/v) FBS and 10 μg/ml ciprofloxacin. The assay medium (II) is based on medium (I) with freshly added 40 ng/ml doxycycline, and 400-600 μM D-luciferin potassium salt (Biosynth). 384-well (Corning 3570, white, flat-bottom, tissue-culture treated) and 6-well plates were first loaded with 20 μl and 1.6 ml of medium (II) for each well respectively. ReBiL cells were trypsinized and cell numbers were determined by Cellometer Auto T4. The required number of cells were collected, centrifuged (200 rcf, 5 min at room temperature) to remove supernatant, and resuspended to 250 cells/μl with medium (I). Then, either 20 μl or 1.6 ml of ReBiL cells were seeded into each well of a 384-well plate (5,000 cells per well) or each well of a 6-well plate (400,000 cells per well), respectively. The final concentration of each component was 1x medium DMEM/F12 (phenol-red free), 10% (v/v) FBS, 10 μg/ml ciprofloxacin, 20 ng/ml doxycycline, and 200-300 μM D-luciferin. The 384-well plate was sealed with MicroAmp Optical Adhesive Film (Thermo Fisher) and luminescence was read in a Tecan M200 microplate reader (integration time 2 sec, 15 min per cycle for a total of 24 hr at 37 °C). The 6-well plate was incubated at 37 °C incubator with 7% CO_2_. Every ReBiL experiment contains a positive control (nl-p53/cl-Mdm2 ReBiL line) and a negative control (nl/cl ReBiL line). Upon termination of the ReBiL assay (24 hr), the Optical Adhesive Film was removed and cell viability was determined using CellTiter-Glo (Promega, 1:5 diluted with PBS, 35 μl per 384-well). To quantify the ½luc fusion proteins, ReBiL cells from 6-well plates were harvested with RIPA lysis buffer (50 mM Tris-HCl pH 8.0, 150 mM NaCl, 0.25% (w/v) deoxycholic acid, 1% (v/v) IGEPAL CA-630, 1 mM EDTA, 2 mM Na_3_VO_4_, 20 mM NaF, and 1x cOmplete Protease Inhibitor cocktail (Roche)) after 24 hrs. The clear lysate was mixed with LDS sample buffer (Thermo Fisher) with 10% (v/v) beta-mercaptoethanol and denatured at 70 °C for 5 min. The denatured lysates were separated by 10% SDS-PAGE and transferred to PVDF membranes (Millipore No. IPFL00010) as described previously[1]. The PVDF membrane was then probed with the anti-HA and anti-Actin antibodies (see Supplemental Table 2) in Odyssey PBS Blocking Buffer (1:1 diluted with PBS) with 0.05% Tween-20 at 4 °C overnight. After incubation with secondary antibodies that are conjugated with Alexa Flour 680 (Thermo Fisher) and IRDye 800 (LI-COR), the membrane was scanned in the LI-COR Odyssey Imaging System (Odyssey Application Software 3.0) using the following parameters: Resolution 84 μm, Quality high, Focus Offset 0.0 mm, Intensity 700 5.0 and 800 5.0. The relative band intensities of [½luc Least], and the actin loading control [loading ctrl] (Extended Data Fig. 2b, c) were determined and exported to Microsoft Excel using Image Studio Software (version 2.1.10, LI-COR) or Image Studio Lite (version 5.2.5, LI-COR). The [½luc Least]/[Actin] was then calculated. The raw luminescent data collected by the Tecan luminometer were imported to Prism 6 (GraphPad). More cells generated higher luminescent signals. This cell number effect was first corrected by calculating the percentage of [Raw ReBiL Luminescence]/[CellTiter Glo Luminescence]. The [Raw ReBiL Luminescence] was the luminescent reading at the 24-hr time point and the [Cell # by CellTiter Glo] was the luminescent reading from CellTiter-Glo cell viability assay performed at the end of ReBiL kinetic assay. Then the ReBiL2.0 score was calculated by dividing the value of ([Raw ReBiL Luminescence]/[CellTiter Glo Luminescence]) by the corresponding ([½luc Least]/[Actin]) (Extended Data Fig. 1d). For each experiment, the ReBiL2.0 scores of nl-p53/cl-Mdm2 and nl/cl were normalized to 100% and 0% respectively. The results are reported as bar charts with the X-axis representing the percent of the p53/Mdm2 interaction.

### KRAS downstream signaling assay

U2OS KRAS ReBiL cells were seeded in a 6-well plate (350,000 - 400,000 cells per well) in DMEM 10% FBS medium with or without 20 ng/ml doxycycline to induce expression of ½luc fusion proteins for 24 hr. The cells were harvested by RIPA lysis buffer and the quantitative western blotting analysis was performed the same as in the ReBiL2.0 assay. For epidermal growth factor (EGF) stimulation experiment, U2OS KRAS ReBiL cells were seeded in 6-well plates (60~70% confluency) and treated with or without 20 ng/ml doxycycline for 24~36 hr. Cells were then washed with PBS and starved in serum free DMEM overnight in the presence or absence of 20 ng/ml doxycycline. Cells were then acutely stimulated with EGF 10 ng/ml (Stemcell Technologies Cat#78006) for 10 min and harvested with RIPA lysis buffer. The quantitative western blotting analysis was performed as described above. The level of phospho-protein was reported as the ratio of the band intensities between the band recognized by the phospho-specific antibody (e.g. [pAKT]) and that recognized by the standard antibody (e.g. [AKT]) on the same western blot using different wave length channels on the LI-COR Image System. The value of phospho-protein from cells without doxycycline was set at 100%.

### Syto60 cell growth assay

Syto60 is a cell-permeant red-fluorescence dye that stains nucleic acid. The intensity of Syto60 fluorescence stain is thus proportional to the cell numbers. U2OS cells carrying TRE-DD-RBD-CAAXK and TRE-DD-RBD were seeded in a 96-well plate (1,000 cells and 200 μl medium (DMEM 10% FBS 10 μg/ml ciprofloxacin, 200 μg/ml G418) per well) with indicated concentration of doxycycline at Day 0. At the times indicated, cells were fixed with 100 μl 10% Buffered Formalin (PROTOCOL 67-56-1) for 10 min at room temperature, washed twice with PBS, and once with TBS (20 mM Tris 150 mM NaCl pH 7.4). Cells were then stained with 1 μM Syto60 dye (Thermo Fisher S11342 prepared in TBS) for 10 min in the dark. Cells were washed three times with TBS. The intensity of Syto60 fluorescence was quantified with the LI-COR Odyssey system.

---

**Supplemental Table.**
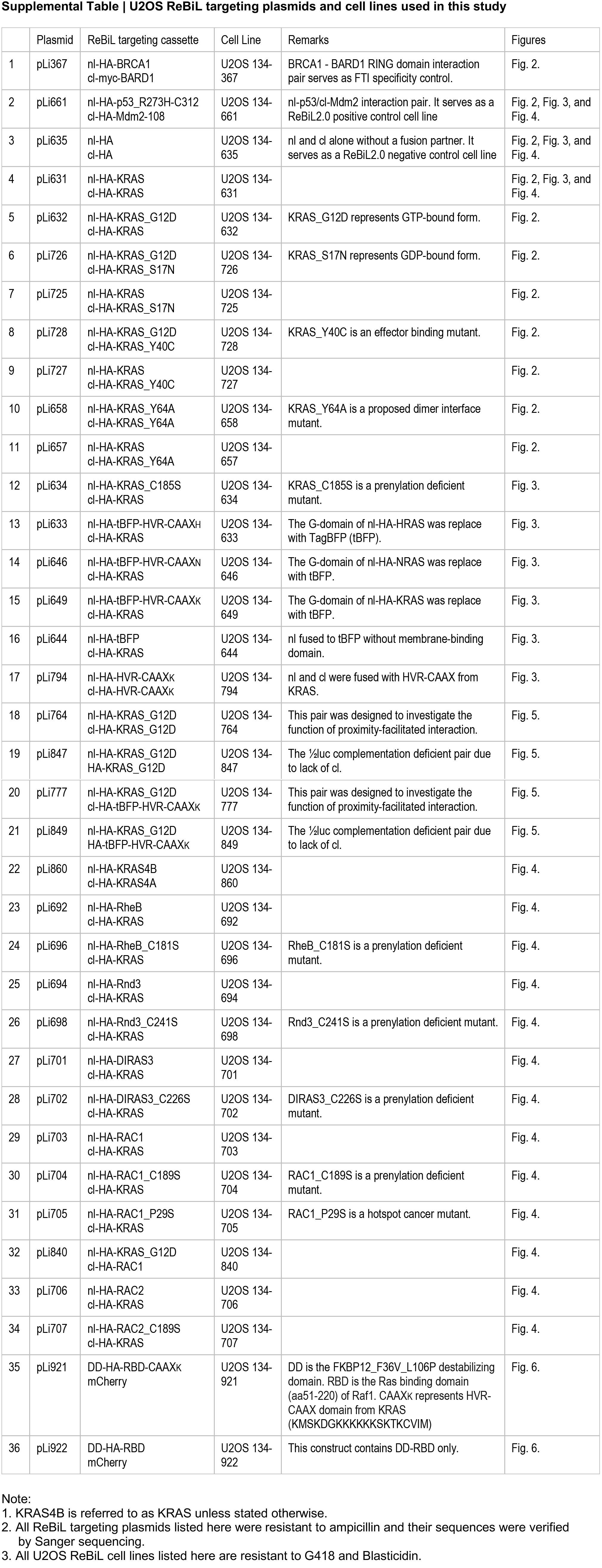
U2OS ReBiL targeting plasmids and cell lines used in this study.

